# A *Xenopus laevis* Model for Systemic Fungal Infections

**DOI:** 10.64898/2026.11.09.765531

**Authors:** Natsuki Atsuta, Ayano Kuriu, Kazuya Ishikawa, Kazuyuki Furuta, Hisao Moriya, Hironobu Nakayama, Chikara Kaito

**Affiliations:** Graduate School of Medicine, Dentistry and Pharmaceutical Sciences, Okayama University, Okayama, 700-8530, Japan; Faculty of Environmental, Life, Natural Science and Technology, Okayama University, Okayama, Japan; Faculty of Pharmaceutical Sciences, Suzuka University of Medical Science, Suzuka 513-8670, Japan

## Abstract

*Xenopus laevis* possesses both innate and adaptive immune systems, and we previously demonstrated its utility as an infection model for human bacterial pathogens. In this study, we investigated the utility of *X. laevis* as a model for fungal infections. The human-pathogenic fungi *Candida albicans* and *Aspergillus fumigatus* exhibited greater lethality in *X. laevis* than the nonpathogenic fungi *Saccharomyces cerevisiae* and *Schizosaccharomyces pombe*. In frogs infected with *C. albicans*, viable fungal cells were detected in the blood and multiple organs early after infection, and gross white lesions and fungal hyphae were observed in the kidneys. *C. albicans* infection also increased renal expression of the proinflammatory cytokine gene*s tnf and il1b*, as well as the tissue injury-associated gene *clu.* Treatment with the antifungal agents amphotericin B and micafungin significantly improved the survival of *C. albicans*-infected frogs. Furthermore, a homozygous deletion mutant of *UME6*, a gene associated with *C. albicans* virulence, exhibited reduced lethality compared with the wild-type strain. These findings establish *X. laevis* as a vertebrate model of systemic fungal infection that enables the evaluation of fungal virulence, host responses, and the therapeutic efficacy of antifungal agents.

## Introduction

Fungal infections are an important global public health concern, causing severe disease particularly in immunocompromised and critically ill patients. *Candida albicans*, *Aspergillus fumigatus*, and *Cryptococcus neoformans* are classified in the critical priority group of the World Health Organization (WHO) Fungal Priority Pathogens List (1). Invasive fungal infections impose a substantial global burden of morbidity and mortality, with mortality rates remaining high even among patients receiving antifungal therapy (2). Moreover, the limited number of available antifungal drug classes and the emergence of antifungal resistance pose major challenges to effective treatment (1). Therefore, a better understanding of fungal pathogenesis and the development of new antifungal agents are needed.

Animal infection models are essential for achieving these goals, as they enable the investigation of fungal virulence and host responses, as well as the evaluation of antifungal efficacy in vivo. Mammalian models, particularly mouse models, have been widely used to study infections caused by human fungal pathogens such as *C. albicans*. For example, the mouse model of systemic candidiasis has contributed substantially to our understanding of host–fungus interactions and the evaluation of antifungal therapies (3). However, mammalian models have practical limitations for experiments requiring large numbers of animals because of ethical and cost considerations. Therefore, the development of complementary infection models is important, particularly from the perspective of the 3Rs (Replacement, Reduction, and Refinement) in animal research (4).

Accordingly, several nonmammalian hosts, including insects and fish, have been developed as models for studying fungal pathogenesis. Invertebrate models such as *Galleria mellonella*, *Drosophila melanogaster*, the silkworm *Bombyx mori*, and the two-spotted cricket *Gryllus bimaculatus* are easy to handle and amenable to experiments involving relatively large numbers of animals and have been used to evaluate fungal virulence factors and antifungal compounds (5–8). In particular, silkworm infection models have been established for several human fungal pathogens and have been used for both virulence studies and antifungal drug evaluation (7, 9–18). Moreover, an *A. fumigatus*-infected silkworm model was successfully used to identify a novel antifungal compound (11). The two-spotted cricket has also been established as an infection model for several human fungal pathogens (5). However, these invertebrate hosts lack an adaptive immune system. The zebrafish, a vertebrate model, offers distinct advantages, particularly the use of transparent larvae for high-throughput in vivo imaging of fungal infections (19–28). Adult zebrafish possess both innate and adaptive immune systems, whereas larvae used in many infection studies lack a fully developed adaptive immune system (29, 30). In addition, zebrafish differ from mammals in several aspects of organ anatomy and physiology, which may limit their ability to reproduce certain aspects of organ-specific pathology associated with systemic fungal infections (31). Thus, additional vertebrate infection models that complement the advantages and limitations of existing models may broaden the experimental approaches available for investigating fungal pathogenesis and host responses.

The African clawed frog, *Xenopus laevis*, is a vertebrate model that has been widely used in developmental biology and immunology (32). *X. laevis* possesses well-developed innate and adaptive immune systems, including B and T lymphocytes, and many components of its immune system are conserved with those of mammals (32, 33). Its relatively large body size also facilitates the collection and analysis of individual organs and blood samples. We previously demonstrated that *X. laevis* is susceptible to infection with human bacterial pathogens, including *Staphylococcus aureus*, *Pseudomonas aeruginosa*, and *Listeria monocytogenes* (34). Mortality caused by these bacterial pathogens was reduced by treatment with appropriate antibacterial agents, and bacterial mutants with attenuated virulence in mammalian hosts also exhibited reduced lethality in *X. laevis* (34). These findings indicate that *X. laevis* can serve as an infection model for evaluating both microbial virulence and antimicrobial efficacy. However, whether *X. laevis* can serve as a model of systemic infection with human fungal pathogens and whether this model can be used to evaluate fungal virulence factors and antifungal therapies remain unknown.

In this study, we investigated the utility of *X. laevis* as a model for systemic fungal infection. We compared the lethality of the human fungal pathogens *C. albicans*, *A. fumigatus*, and *C. neoformans* with that of nonpathogenic fungi and characterized the dissemination of *C. albicans* and the associated inflammatory responses, tissue injury, and hyphal formation in the kidney. We further evaluated the therapeutic efficacy of amphotericin B and micafungin and examined the effect of deleting *UME6*, which encodes a transcriptional regulator required for hyphal extension and virulence in *C. albicans* (35, 36). Together, these experiments were designed to determine whether *X. laevis* can provide a vertebrate model for investigating systemic fungal pathogenesis and evaluating fungal virulence factors and antifungal therapies.

## Results

### Intraperitoneal inoculation of *Candida albicans* causes lethal infection in *Xenopus laevis*

To determine whether *X. laevis* is susceptible to infection with a human fungal pathogen, frogs were intraperitoneally inoculated with *Candida albicans*. Frogs that died within 24 h after inoculation exhibited marked reddish discoloration of the abdominal region (**Fig. 1A**).

**Fig. 1.**
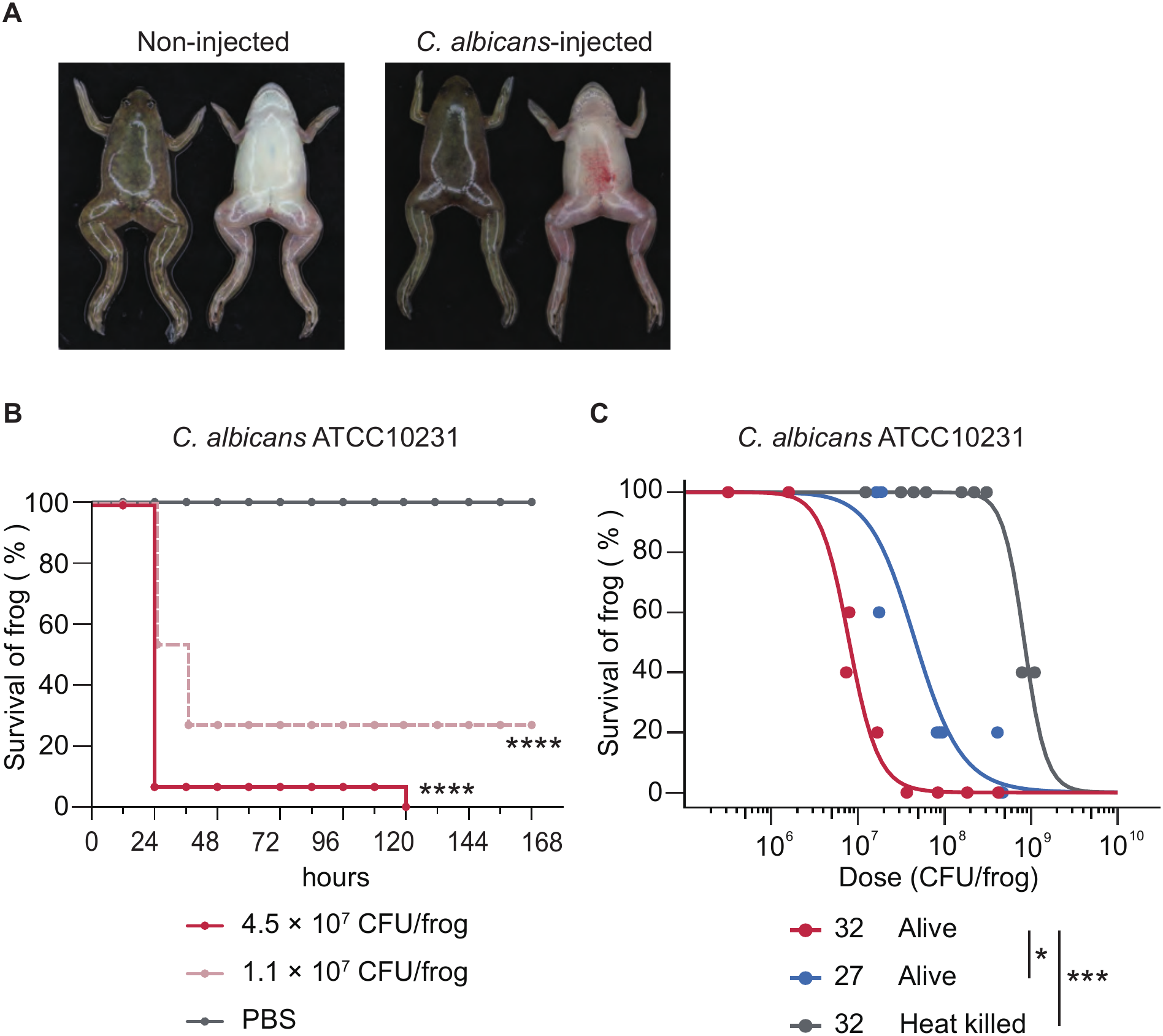
Intraperitoneal inoculation with *Candida albicans* causes lethal infection in Xenopus laevis. (A) Representative images of frogs at 24 h after intraperitoneal inoculation with PBS (left) or *C. albicans* ATCC 10231 (4.5 × 10⁷ CFU/frog; right). The *C. albicans*-inoculated frog had died by 24 h, whereas the Non-injected frog remained alive. (B) Survival of frogs following intraperitoneal inoculation with live *C. albicans* ATCC 10231 at 4.5 × 10⁷ or 1.125 × 10⁷ CFU/frog. Five frogs were used per group in each experiment, and data from three independent experiments were pooled (n = 15). Survival curves were compared with that of the PBS-injected group using the log-rank (Mantel-Cox) test. ****, *P* < 0.0001. (C) Dose-response analysis of frogs inoculated with different doses of live or heat-killed *C. albicans* ATCC 10231. Frogs inoculated with live *C. albicans* were maintained at either 27°C or 32°C. Heat-killed fungal cells were prepared by autoclaving live *C. albicans* at 121°C for 20 min. Survival was determined at 72 h after inoculation. Three fungal doses were tested in each experiment, with five frogs used per dose. Three independent experiments were performed, and dose-response curves were generated by logistic regression. Differences between the dose-response curves at 27°C and 32°C and between live and heat-killed *C. albicans* were evaluated by likelihood ratio tests. *, *P* < 0.05; ***, *P* < 0.001.

We next examined the time course and dose dependence of *C. albicans*-induced mortality. Frogs were intraperitoneally inoculated with different doses of *C. albicans* and monitored for survival. Deaths were observed beginning at 24 h after inoculation, and survival decreased with increasing inoculum size (**Fig. 1B**), demonstrating that *C. albicans* causes dose-dependent lethal infection in *X. laevis*.

We then examined whether the lethality of *C. albicans* required viable fungal cells by comparing the 50% lethal doses (LD50) of live and heat-killed *C. albicans*. The LD50 of live *C. albicans* was 8.0 × 10⁶ CFU/frog, whereas that of heat-killed *C. albicans* was >8.4 × 10⁸ CFU/frog (**Fig. 1C**). We also examined the effect of incubation temperature on *C. albicans* lethality. At 27°C, the LD50 of live *C. albicans* at 72 h after inoculation was 4.8 × 10⁷ CFU/frog, more than fivefold higher than that at 32°C (**Fig. 1C**). These results indicate that the lethality of *C. albicans* in *X. laevis* depends on fungal viability and is greater at 32°C than at 27°C.

### Multiple human fungal pathogens cause lethal infection in *Xenopus laevis*

In addition to *C. albicans* ATCC 10231, we examined whether *C. albicans* SC5314 and the human fungal pathogens *Aspergillus fumigatus* and *Cryptococcus neoformans* caused lethal infection in *X. laevis*. For *A. fumigatus* and *C. neoformans*, the lethality of live and heat-killed fungal cells was also compared. Similar to *C. albicans* ATCC 10231 (**Fig. 1C**), intraperitoneal inoculation with live *C. albicans* SC5314, *A. fumigatus*, or *C. neoformans* resulted in dose-dependent decreases in frog survival (**Fig. 2A–C**). The LD50 values calculated from the survival curves were 1.4 × 10⁶ CFU/frog for *C. albicans* SC5314, 5.1 × 10⁷ CFU/frog for *A. fumigatus*, and 1.8 × 10⁸ CFU/frog for *C. neoformans*. In contrast, frogs inoculated with heat-killed *A. fumigatus* or *C. neoformans* showed higher survival than those inoculated with the corresponding live fungi, and LD50 values could not be determined within the range of inoculum sizes tested (**Fig. 2B and C**). These results indicate that fungal viability contributes substantially to the lethality of *A. fumigatus* and *C. neoformans* in *X. laevis*.

**Fig. 2.**
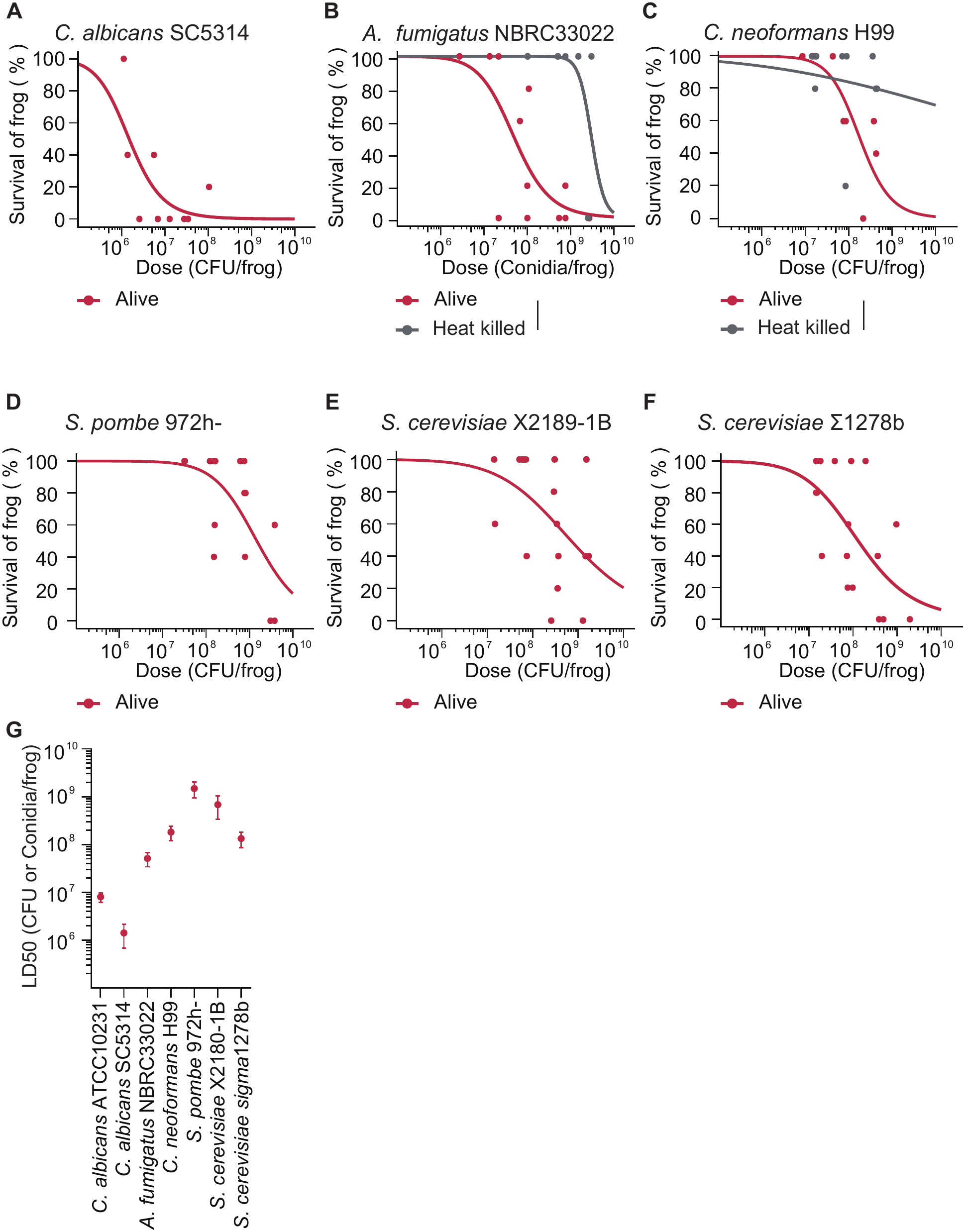
Lethality of different fungal species and strains in *Xenopus laevis*. (A–F) Dose-response analysis of frogs inoculated with different doses of *Candida albicans* SC5314 (A), *Aspergillus fumigatus* NBRC 33022 (B), *Cryptococcus neoformans* H99 (C), *Schizosaccharomyces pombe* 972h^−^ (D), *Saccharomyces cerevisiae* X2180-1B (E), or *S. cerevisiae* Σ1278b (F). Three fungal doses were tested in each experiment, with five frogs used per dose. Following inoculation, frogs were maintained at 32°C, and survival was determined at 72 h after inoculation. Three independent experiments were performed, and dose-response curves were generated by logistic regression. Where applicable, differences between live and heat-killed fungi were evaluated by likelihood ratio tests. *, *P* < 0.05; ***, *P* < 0.001. (G) LD50 values calculated from the dose-response curves for live fungi shown in Fig. 1C and panels A–F. Error bars indicate standard errors.

We next examined the lethality of the nonpathogenic fungi *Schizosaccharomyces pombe*, *Saccharomyces cerevisiae* X2180-1B, and *S. cerevisiae* Σ1278b. Although high inoculum sizes of all three fungi caused frog mortality, the extent of lethality differed among the species and strains tested (**Fig. 2D–F**). The LD50 values were 1.5 × 10⁹ CFU/frog for *S. pombe*, 6.8 × 10⁸ CFU/frog for *S. cerevisiae* X2180-1B, and 1.3 × 10⁸ CFU/frog for *S. cerevisiae* Σ1278b.

Comparison of the LD50 values showed that *C. albicans* and *A. fumigatus* caused lethal infection at lower inoculum sizes than *S. pombe* and *S. cerevisiae* X2180-1B (**Fig. 2G**). In contrast, *C. neoformans* exhibited lower lethality than *C. albicans* and *A. fumigatus*, with an LD50 similar to that of *S. cerevisiae* Σ1278b. Thus, *X. laevis* is susceptible to multiple human fungal pathogens, although the degree of lethality varies considerably among fungal species and strains.

### *Candida albicans* disseminates systemically following intraperitoneal inoculation in *Xenopus laevis*

To characterize the dissemination of *C. albicans* following intraperitoneal inoculation, *X. laevis* frogs were inoculated with *C. albicans* ATCC 10231 and dissected at 0.5, 6, 12, 18, and 24 h after inoculation. Viable fungal burdens in the blood, liver, kidney, and brain were determined by CFU enumeration. *C. albicans* was detected in the blood and all organs examined as early as 0.5 h after inoculation. Fungal burdens remained relatively stable in the blood, liver, and brain for up to 24 h, whereas the fungal burden in the kidney increased over time (**Fig. 3A**).

**Fig. 3.**
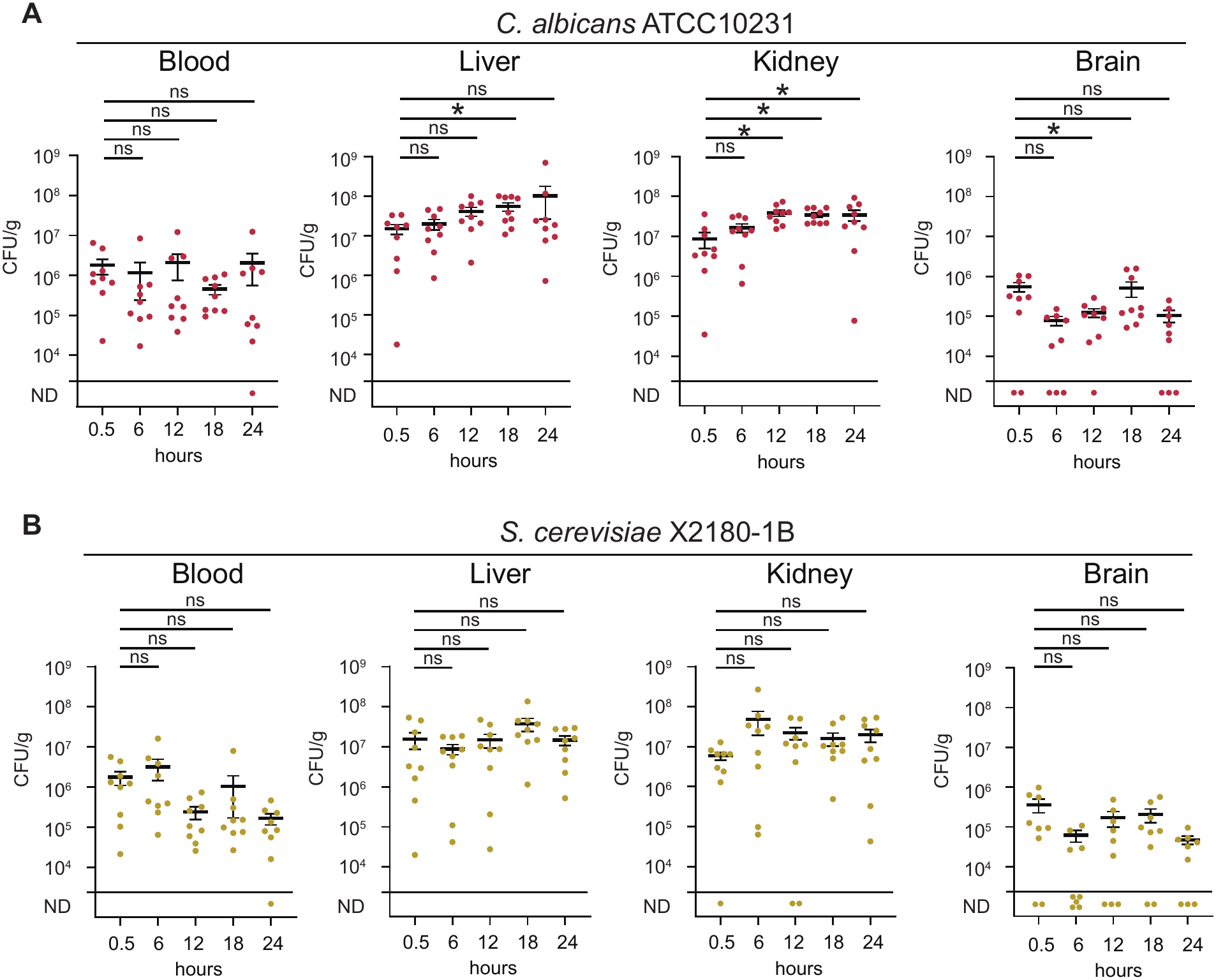
*Candida albicans* disseminates systemically following intraperitoneal inoculation in *Xenopus laevis*. (A) Frogs were intraperitoneally inoculated with *C. albicans* ATCC 10231 (4.5 × 10⁷ CFU/frog) and maintained at 32°C. At 0.5, 6, 12, 18, and 24 h after inoculation, surviving frogs were dissected, and viable fungal burdens in the blood, liver, kidneys, and brain were determined by CFU enumeration. Fungal burdens were expressed as CFU/mL of blood or CFU/g of tissue. (B) Frogs were intraperitoneally inoculated with *S. cerevisiae* X2180-1B (4.5 × 10⁷ CFU/frog), and fungal burdens in the blood, liver, kidneys, and brain were determined at the indicated time points as described for panel A. Three frogs were used at each time point in each experiment, and three independent experiments were performed (n = 9). For each organ or blood, fungal burdens at each time point were compared with those at 0.5 h using one-way ANOVA followed by Dunnett’s multiple comparisons test. ND, below the detection limit.

We next examined whether a similar distribution occurred with the nonpathogenic fungus *S. cerevisiae*. Frogs were intraperitoneally inoculated with *S. cerevisiae* X2180-1B, and viable fungal burdens in the blood and organs were determined as described above. *S. cerevisiae* was also detected in the blood and all organs examined (**Fig. 3B**). At 24 h after inoculation, the fungal burdens of *S. cerevisiae* in the kidney and brain were comparable to those of *C. albicans*, whereas those in the blood and liver were approximately 10-fold lower. In addition, *S. cerevisiae* burdens remained relatively stable in all organs throughout the observation period, with no marked increase (**Fig. 3B**).

These results demonstrate that intraperitoneally inoculated *C. albicans* rapidly disseminates throughout the body and persists in the blood and multiple organs for at least 24 h, with a progressive increase in fungal burden in the kidney. Although *S. cerevisiae* also disseminated systemically, its burdens in the blood and liver were lower than those of *C. albicans*, suggesting a reduced capacity to persist or proliferate in these tissues.

### *Candida albicans* induces proinflammatory cytokine gene expression in the kidneys of *Xenopus laevis*

Because the fungal burden of *C. albicans* increased in the kidney after infection, we next examined the renal inflammatory response. Frogs were intraperitoneally inoculated with *C. albicans* ATCC 10231 or *S. cerevisiae* X2180-1B, and kidneys were collected at 24 h after inoculation. Total RNA was extracted from the kidneys, and the expression levels of the proinflammatory cytokine genes *tnf* and *il1b* were determined by RT-qPCR. In frogs inoculated with *C. albicans*, the expression levels of both *tnf* and *il1b* were significantly increased compared with those in PBS-injected controls (**Fig. 4A and B**). In contrast, *S. cerevisiae* inoculation significantly increased *tnf* expression but did not significantly increase *Il1b* expression compared with PBS injection (**Fig. 4C and D**). These results indicate that *C. albicans* infection induces the expression of proinflammatory cytokine genes in the *X. laevis* kidney, whereas *S. cerevisiae* induces a more limited cytokine response under these experimental conditions.

**Fig. 4.**
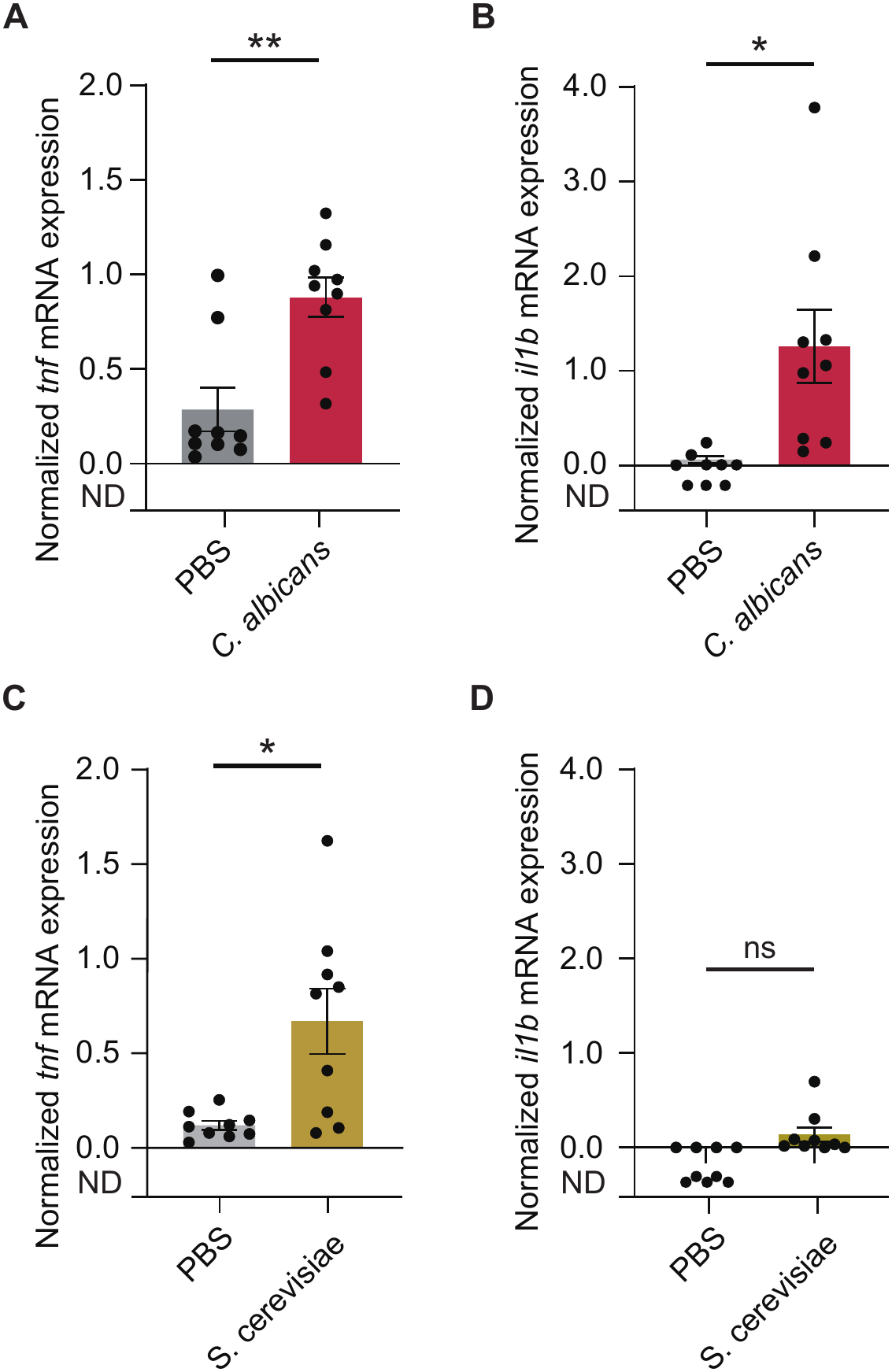
*Candida albicans* infection induces proinflammatory cytokine gene expression in the kidneys of *Xenopus laevis*. (A and C) Relative expression of *tnf* gene in the kidneys of frogs inoculated with *C. albicans* (A) or *S. cerevisiae* (C). (B and D) Relative expression of *Il1b* gene in the kidneys of frogs inoculated with *C. albicans* (B) or *S. cerevisiae* (D). Frogs were intraperitoneally inoculated with *C. albicans* ATCC 10231 (4.5 × 10⁷ CFU/frog), *S. cerevisiae* X2180-1B (4.5 × 10⁷ CFU/frog), or PBS. At 24 h after inoculation, kidneys were collected from surviving frogs, and total RNA was extracted. Relative transcript levels were determined by RT-qPCR and normalized to β-actin. Three frogs were analyzed per group in each experiment, and three independent experiments were performed (n = 9). Fungal-inoculated groups were compared with the PBS-injected control group using unpaired *t* tests with Welch’s correction. ns, *P* > 0.05; **, *P* < 0.01; ***, *P* < 0.001. ND, not detected.

### *Candida albicans* infection increases the expression of a tissue injury-associated gene in the kidney

Because *C. albicans* infection resulted in an increased fungal burden and an inflammatory response in the kidney, we next examined whether the infection was associated with renal tissue injury. We analyzed the expression of *clu*, which encodes clusterin, a stress-responsive protein whose expression is induced following cellular and tissue injury in mammals (37). In adult *X. laevis*, *clu* is expressed at a low level in the kidney (35). RT-qPCR analysis showed that renal *clu* expression was significantly increased in *C. albicans*-infected frogs compared with PBS-injected controls (**Fig. 5**). These results demonstrate that *C. albicans* infection induces renal *clu* expression in *X. laevis*.

**Fig. 5.**
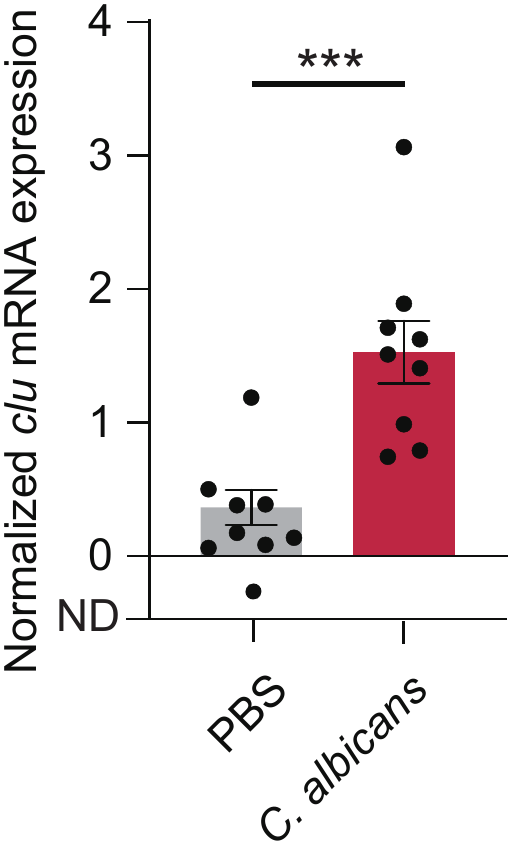
*Candida albicans* infection increases the expression of a tissue injury-associated gene in the kidneys of *Xenopus laevis*. Relative expression of Clu (Clusterin) gene in the kidneys of frogs inoculated with *C. albicans*. Frogs were intraperitoneally inoculated with *C. albicans* ATCC 10231 (4.5 × 10⁷ CFU/frog) or PBS. At 24 h after inoculation, kidneys were collected from surviving frogs, and total RNA was extracted. The relative transcript level of *clu* was determined by RT-qPCR and normalized to β-actin. Three frogs were analyzed per group in each experiment, and three independent experiments were performed (n = 9). The *C. albicans*-inoculated group was compared with the PBS-injected control group using an unpaired *t* test with Welch’s correction. ns, *P* > 0.05; **, *P* < 0.01; ***, *P* < 0.001.

### *Candida albicans* infection causes gross white lesions in the kidneys of *Xenopus laevis*

Because *C. albicans* infection induced changes in the expression of proinflammatory cytokine and tissue injury-associated genes in the kidney, we next examined whether gross pathological changes developed in the kidneys. Compared with PBS-injected controls, frogs intraperitoneally inoculated with *C. albicans* ATCC 10231 exhibited discoloration of the abdominal organs and multiple white lesions in the kidneys (**Fig. 6A and B**). In contrast, similar white lesions were not observed in the kidneys of frogs inoculated with PBS or *S. cerevisiae* X2180-1B (**Fig. 6A and 6C**). These results demonstrate that *C. albicans* infection causes characteristic gross lesions in the kidneys of *X. laevis*.

**Fig. 6.**
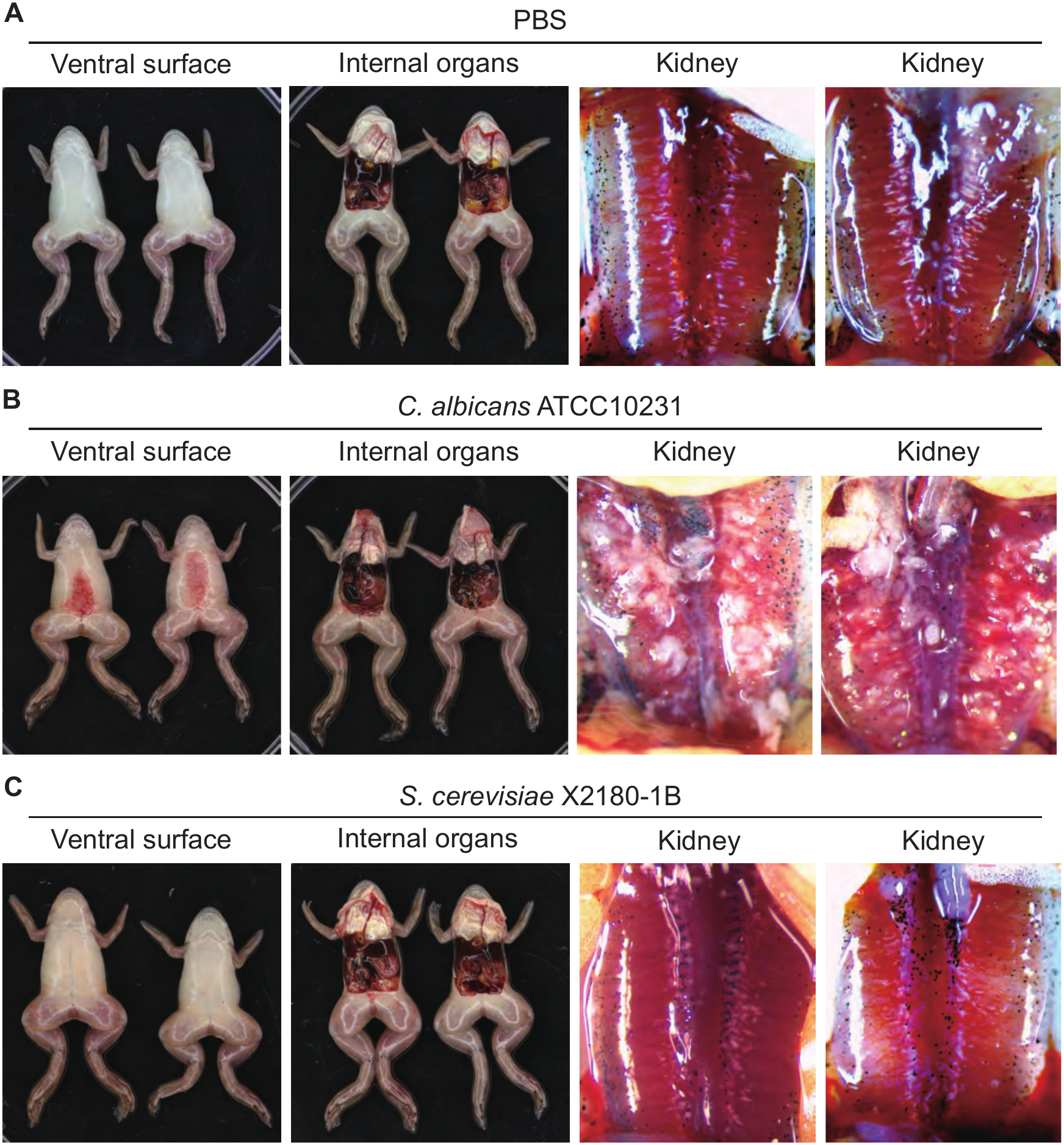
*Candida albicans* infection causes gross white lesions in the kidneys of Xenopus laevis. Frogs were intraperitoneally inoculated with PBS (A), *C. albicans* ATCC 10231 (4.5 × 10⁷ CFU/frog) (B), or *S. cerevisiae* X2180-1B (4.5 × 10⁷ CFU/frog) (C) and maintained at 32°C. At 24 h after inoculation, surviving frogs were anesthetized and photographed. Images show the ventral view of the whole frog, the abdominal cavity after dissection, and a magnified view of the kidneys in situ. Representative images from two frogs in each group are shown.

### *Candida albicans* forms hyphae in the kidneys of *Xenopus laevis*

Because gross white lesions were observed in the kidneys following *C. albicans* infection, we next examined fungal morphology and histopathological changes in the kidney. Kidney sections collected 24 h after infection were stained with hematoxylin and eosin (H&E) and periodic acid-Schiff (PAS). In frogs infected with either *C. albicans* ATCC 10231 or SC5314, prominent PAS-positive hyphae were observed in the kidney tissue, whereas no fungal structures were detected in the kidneys of PBS-injected controls (**Fig. 7**). In contrast, only a small number of yeast-like cells were detected locally in the kidney tissue of frogs inoculated with *S. cerevisiae* X2180-1B (**Fig. 7**). These results indicate that *C. albicans* disseminates to the kidney following intraperitoneal inoculation and forms hyphae within the kidney tissue.

**Fig. 7.**
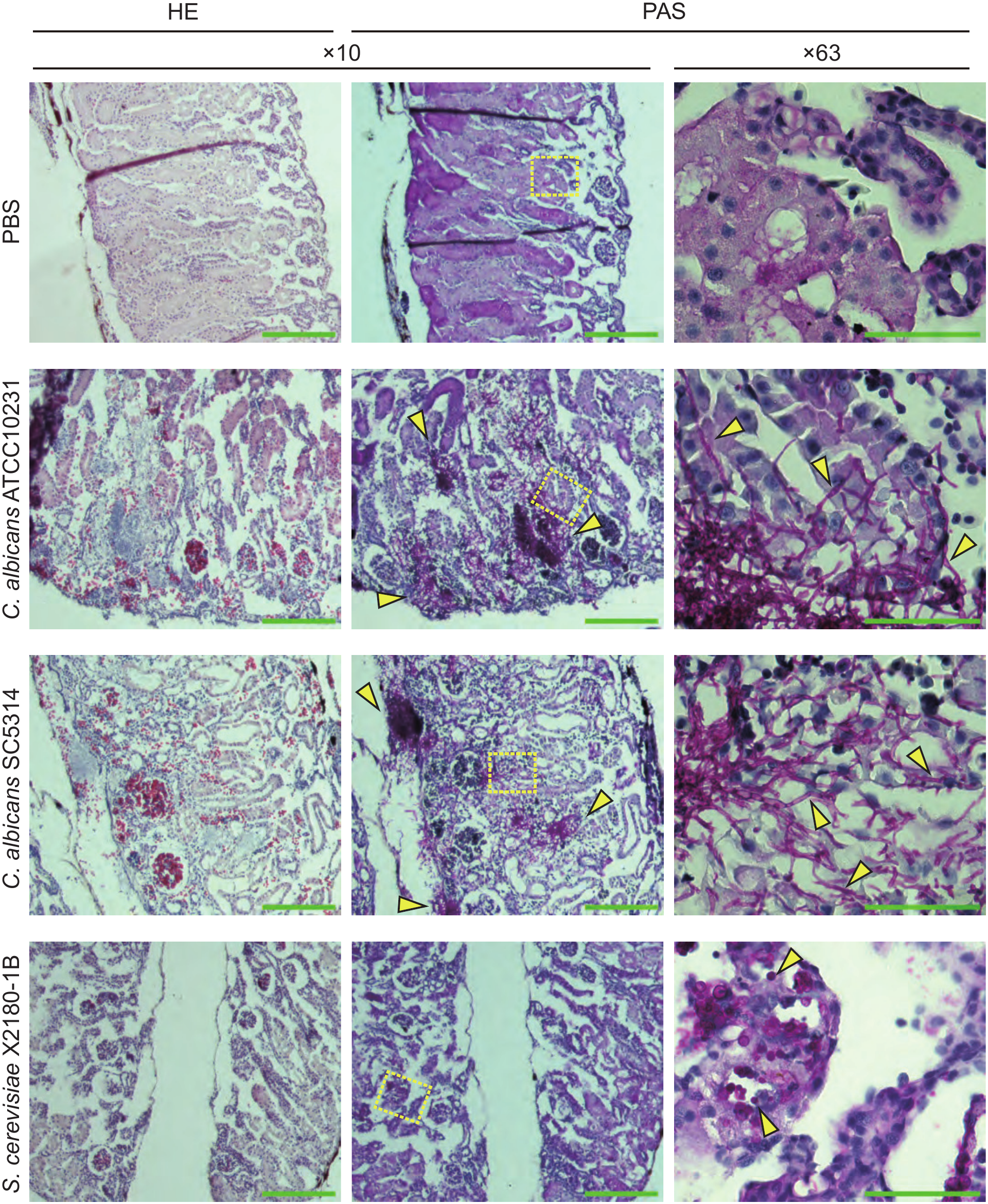
*Candida albicans* forms hyphae in the kidneys of *Xenopus laevis*. Frogs were intraperitoneally inoculated with *C. albicans* ATCC 10231, *C. albicans* SC5314, *S. cerevisiae* X2180-1B (4.5 × 10⁷ CFU/frog each), or PBS and maintained at 32°C. At 24 h after inoculation, kidneys were collected from surviving frogs, fixed, and embedded in paraffin. Tissue sections were stained with hematoxylin and eosin (H&E) or periodic acid–Schiff (PAS) and examined using 10× or 63× objective lenses. In H&E-stained sections, nuclei appear blue-purple and the cytoplasm appears pink. In PAS-stained sections, polysaccharide-rich structures, including fungal cells and hyphae, appear magenta, whereas hematoxylin-counterstained nuclei appear blue-purple. Yellow boxes in the 10× images indicate the regions shown at higher magnification in the corresponding 63× images. Yellow arrows indicate fungal cells or hyphae. Scale bars, 200 μm for images obtained with the 10× objective and 50 μm for images obtained with the 63× objective.

### Antifungal treatment improves survival of *Candida albicans*-infected *Xenopus laevis*

To determine whether the *X. laevis* infection model could be used to evaluate the therapeutic efficacy of antifungal agents, we examined the effects of amphotericin B and micafungin on survival following *C. albicans* infection. When amphotericin B or micafungin was administered intraperitoneally immediately after *C. albicans* inoculation, both treatments significantly improved frog survival compared with the respective control groups (**Fig. 8A and B**).

**Fig. 8.**
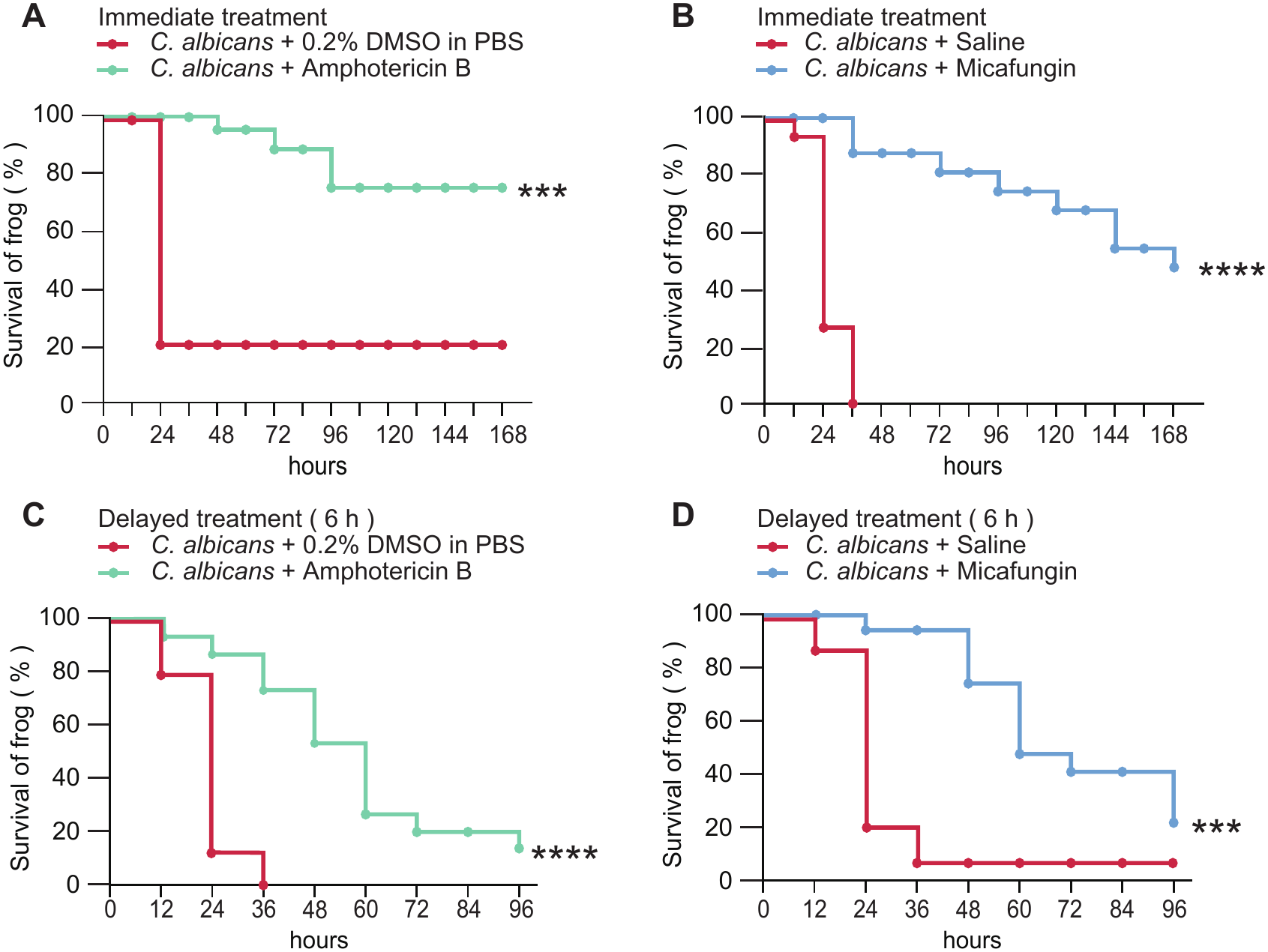
Antifungal treatment improves survival of *Candida albicans*-infected *Xenopus laevis*. (A and B) Frogs were intraperitoneally inoculated with *C. albicans* ATCC 10231 (4.5 × 10⁷ CFU/frog), followed immediately by intraperitoneal administration of amphotericin B (5 μg/frog; approximately 2 mg/kg) or its vehicle (0.2%DMSO in PBS) (A), or micafungin (15 μg/frog; approximately 6 mg/kg) or its vehicle (0.9%NaCl) (B). (C and D) Frogs were intraperitoneally inoculated with *C. albicans* ATCC 10231 (4.5 × 10⁷ CFU/frog), followed at 6 h after inoculation by intraperitoneal administration of amphotericin B (5 μg/frog; approximately 2 mg/kg) or its vehicle (0.2%DMSO in PBS) (C), or micafungin (15 μg/frog; approximately 6 mg/kg) or its vehicle (0.9%NaCl) (D). Survival was monitored at 12-h intervals after antifungal administration. Five frogs were used per group in each experiment, and data from three independent experiments were pooled (n = 15). Survival curves were compared between the antifungal-treated and corresponding vehicle control groups using the log-rank (Mantel-Cox) test. ***, *P* < 0.001; ****, *P* < 0.0001.

We next examined whether antifungal treatment was effective when initiated after systemic dissemination of *C. albicans*. Amphotericin B or micafungin was administered intraperitoneally at 6 h after *C. albicans* inoculation, a time point at which the fungus had already disseminated to multiple organs (**Fig. 3A**). Both treatments significantly improved frog survival compared with the respective control groups (**Fig. 8C and D**). These results demonstrate that amphotericin B and micafungin exert therapeutic effects against *C. albicans* infection in *X. laevis*, even when administered after systemic fungal dissemination. Thus, the *X. laevis* infection model can be used to evaluate the in vivo efficacy of antifungal agents.

### Candida albicans UME6 contributes to virulence in Xenopus laevis

To determine whether the *X. laevis* infection model could be used to evaluate the contribution of fungal genes to virulence, we examined a *C. albicans UME6* deletion mutant. *UME6* encodes a transcriptional regulator that is required for hyphal extension and contributes to *C. albicans* virulence (35). Frogs intraperitoneally inoculated with a homozygous *UME6* deletion mutant (Δ/Δ *ume6*) showed significantly increased survival compared with those inoculated with the wild-type (WT) strain (**Fig. 9**). Both heterozygous UME6 deletion mutants (Δ/+ ume6), generated using different selectable markers, exhibited survival curves intermediate between those of the WT and homozygous deletion strains, although their survival did not differ significantly from that of the WT strain (**Fig. 9**). These results demonstrate that homozygous deletion of *UME6* attenuates the virulence of *C. albicans* in *X. laevis*, indicating that *UME6* contributes to virulence in this infection model. Furthermore, these findings support the utility of the *X. laevis* infection model for evaluating the contributions of fungal genes to virulence.

**Fig. 9.**
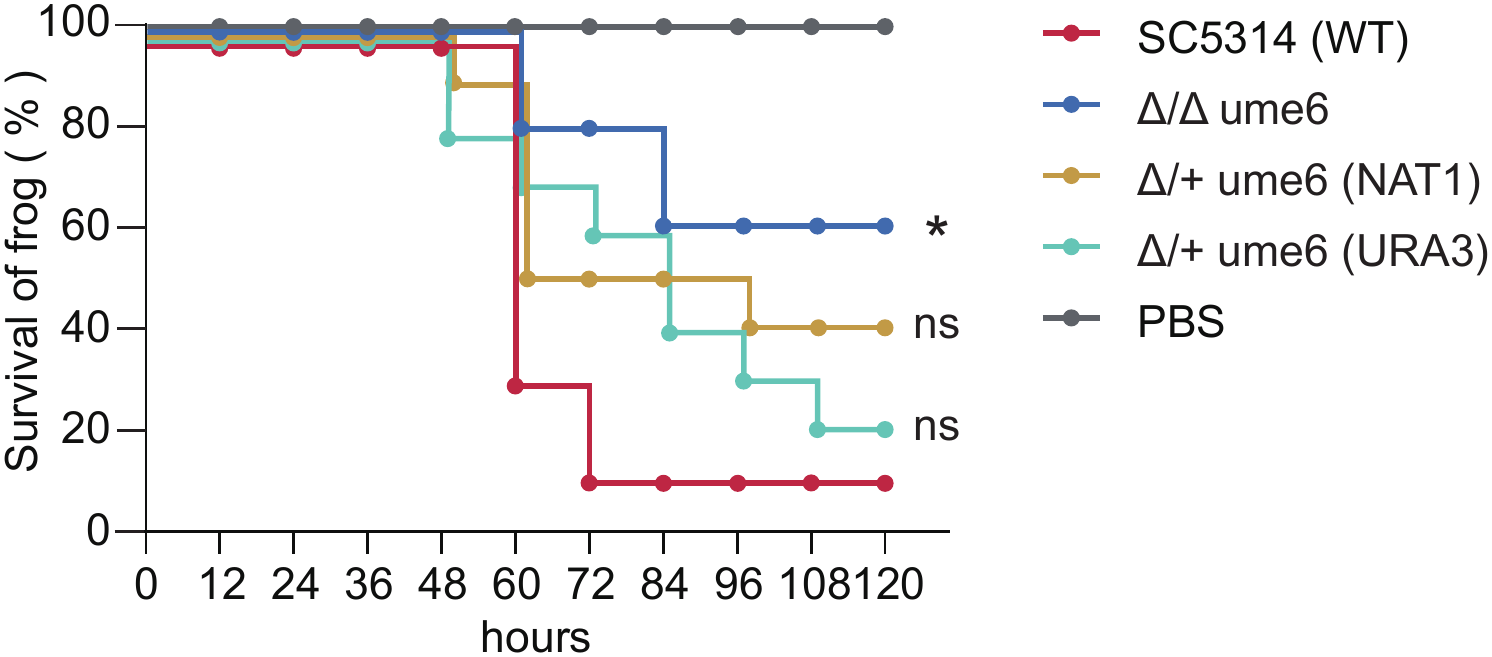
Homozygous deletion of *UME6* attenuates *Candida albicans* virulence in Xenopus laevis. Frogs were intraperitoneally inoculated with *C. albicans* SC5314 (WT), a homozygous *UME6* deletion mutant (Δ/Δ *ume6*), or either of two heterozygous *UME6* deletion mutants [Δ/+ *ume6* (NAT1) and Δ/+ *ume6* (URA3)] at 5.0 × 10⁶ CFU/frog and maintained at 32°C. Five frogs were used per group in each experiment, and data from two independent experiments were pooled (n = 10). Survival curves were compared with that of the WT strain using the log-rank test with Holm-Šídák correction for multiple comparisons. **, *P* < 0.01.

## Discussion

In this study, we demonstrate that the African clawed frog, *Xenopus laevis*, can serve as a vertebrate model for systemic infection with human fungal pathogens. *C. albicans* and *A. fumigatus* exhibited greater lethality than the nonpathogenic fungi tested, and *C. albicans* disseminated to multiple organs following intraperitoneal inoculation, with a progressive increase in fungal burden in the kidney. *C. albicans* infection was also associated with increased expression of proinflammatory cytokine genes and *clu* in the kidney, as well as gross white lesions and hyphal formation. Furthermore, amphotericin B and micafungin improved the survival of infected frogs, whereas homozygous deletion of *UME6* attenuated *C. albicans* virulence. Together, these findings support the utility of *X. laevis* for investigating fungal virulence, host responses and tissue pathology during infection, and the therapeutic efficacy of antifungal agents.

The prominent kidney involvement observed during *C. albicans* infection resembles an important feature of disseminated candidiasis in mammals. In mouse models, *C. albicans* disseminates to multiple organs and subsequently accumulates in the kidney, where fungal proliferation, hyphal formation, and inflammatory responses contribute to tissue damage (38–40). Similarly, in *X. laevis*, we observed a progressive increase in viable *C. albicans* in the kidney, increased expression of *tnf*, *il1b*, and *clu*, and the development of gross white lesions containing fungal hyphae. Thus, despite differences in infection route and body temperature, *C. albicans* infection in *X. laevis* shares several features with mammalian systemic candidiasis, particularly fungal accumulation and associated pathological changes in the kidney. Further characterization of the white renal lesions, including their fungal and host immune cell components, will be important to determine whether they resemble the renal abscesses observed during disseminated candidiasis in mammals.

Hyphal growth is a major virulence-associated property of *C. albicans* that contributes to tissue invasion and damage, and Ume6 is a key transcriptional regulator of hyphal extension (35, 36). In mouse models, loss of *UME6* impairs hyphal extension and attenuates virulence (35). In the present study, *C. albicans* formed prominent hyphae within the *X. laevis* kidney, and homozygous deletion of *UME6* significantly attenuated virulence. These findings indicate that Ume6 also contributes to *C. albicans* virulence in an amphibian host and support the use of this model for evaluating established fungal virulence factors. However, because Ume6 regulates genes involved in processes other than hyphal growth, the reduced virulence of the *UME6* mutant cannot be attributed solely to impaired hyphal formation.

The differences in lethality among the fungal species and strains tested also raise interesting questions regarding the contribution of fungal morphology to virulence in *X. laevis*. Among the nonpathogenic *S. cerevisiae* strains, Σ1278b exhibited greater lethality than X2180-1B. The Σ1278b background is well known for its capacity for pseudohyphal and invasive growth, phenotypes in which the cell surface protein Flo11 plays an important role (41, 42). The greater lethality of Σ1278b is therefore consistent with the possibility that filamentous or invasive growth contributes to fungal virulence in *X. laevis*. However, whether Σ1278b undergoes pseudohyphal growth within the frog remains unknown, and this possibility requires direct examination. In contrast, *C. neoformans* exhibited lower lethality than *C. albicans* and *A. fumigatus*, with an LD50 similar to that of *S. cerevisiae* Σ1278b. Unlike *C. albicans* and *A. fumigatus*, *C. neoformans* generally proliferates as a yeast during mammalian infection and relies on virulence attributes such as its polysaccharide capsule, melanin production, and thermotolerance rather than invasive hyphal growth (43, 44). Moreover, the experimental temperature of 32°C may differentially affect the expression of virulence traits among these fungi. Because cryptococcosis is typically characterized by a more prolonged course of infection, it will also be of interest to determine whether *X. laevis* can be adapted to study persistent or chronic *C. neoformans* infection.

The ability of amphotericin B and micafungin to improve survival when administered not only immediately after inoculation but also 6 h after infection, when *C. albicans* had already disseminated to multiple organs, demonstrates that this model can detect the *in vivo* therapeutic effects of antifungal agents with different mechanisms of action. Further studies examining dose-response relationships and reductions in organ fungal burdens will help establish the utility of this model for evaluating novel antifungal compounds.

Several nonmammalian hosts, including insects and zebrafish, have been used as models of fungal infection (5–7). Zebrafish larvae, in particular, offer the major advantage of optical transparency, enabling high-throughput in vivo imaging of interactions between fungi and innate immune cells. However, the adaptive immune system is not fully developed during the larval stages commonly used for infection studies (45). Adult zebrafish possess both innate and adaptive immunity, but their kidneys also serve as major hematopoietic organs and differ from mammalian kidneys in several anatomical and physiological respects. Adult *X. laevis*, in contrast, possesses both innate and adaptive immune systems (32) and has several histological and functional features of major organs in common with mammals (46). This feature may be advantageous for studying organ-level host–fungus interactions, as *C. albicans*, *A. fumigatus*, and *C. neoformans* cause characteristic pathological changes in mammalian organs, including the kidneys, lungs, and central nervous system (47–49). In addition, the relatively large body size of *X. laevis* allows sufficient samples to be obtained individually from the blood, liver, kidney, and brain, facilitating the combined analysis of organ fungal burdens, gene expression, gross pathology, and histopathology. Thus, whereas zebrafish provide particular advantages for high-throughput *in vivo* imaging, *X. laevis* may provide a complementary vertebrate model for multifaceted analysis of organ-level pathology and host responses during systemic fungal infection.

This study has several limitations. First, the experiments were performed at 32°C, which is below human body temperature and may influence fungal growth and virulence factor expression. Second, the mechanisms underlying the renal lesions were not fully characterized. Third, detailed analyses of dissemination, host responses, and tissue pathology were largely restricted to *C. albicans*. Because *A. fumigatus* and *C. neoformans* are naturally acquired primarily through the respiratory tract (43, 49), future studies should explore alternative routes of infection in *X. laevis* and characterize the resulting tissue pathology. Finally, direct assessment of fungal burdens and filamentous growth in frogs infected with the *C. albicans UME6* mutant or *S. cerevisiae* Σ1278b will be required to more clearly define the relationship between filamentous growth and lethality in this model.

In conclusion, our findings establish *X. laevis* as a vertebrate model for investigating systemic infections caused by human fungal pathogens. This model enables the assessment of systemic fungal dissemination, organ pathology, host responses, antifungal efficacy, and the contributions of fungal genes to virulence. *X. laevis* therefore provides a complementary experimental system to existing mammalian, fish, and invertebrate models for studying fungal pathogenesis and evaluating antifungal therapies.

## Materials and Methods

### Xenopus laevis

Juvenile *Xenopus laevis* (3–4 cm in body length) were purchased from Xenopus Yoshoku Kyozai (Ibaraki, Japan). The frogs had an average body weight of approximately 2.5 g. Frogs were maintained in a temperature-controlled room at 22°C in 13.6-liter tanks (37 cm wide × 22 cm deep × 25 cm high), with 50 frogs per tank. The frogs were fed three times per week, and the tank water was replaced three times per week.

### Preparation of fungal suspensions

The fungal strains used in this study are listed in Table 1. *Candida albicans* and *Cryptococcus neoformans* were cultured aerobically in YPD medium at 30°C for 24 h with shaking. *Saccharomyces cerevisiae* was cultured in YPD medium and *Schizosaccharomyces pombe* in YES medium at 30°C for 48 h with shaking. Fungal cultures were centrifuged at 16,200 × *g* for 10 min, and the resulting pellets were resuspended in phosphate-buffered saline (PBS) and diluted to the desired concentrations.

**Table 1.** Strains used in this study.

| Strain | Parent strain | Genotype or characteristics | Source |
|---|---|---|---|
| *C. albicans* |  |  |  |
| ATCC10231 |  | Serotype A | (60) |
| SC5314 |  | Wild type | (61) |
| CAF2 | SC5314 | *URA3/ura3::imm434* | (58) |
| CAI8 | CAF2 | *ade2::hisG/ade2::hisG ura3::imm434/ura3::imm434* | (58) |
| EGFP | CAI8 | *ade2::hisG/ade2::hisG ura3::imm434/ura3::imm434 ENO1/eno1::P<sub>ENO1</sub>-yGFP3* | This study |
| EGFP-URA3 | EGFP | *ade2::hisG/ade2::hisG URA3/ura3::imm434 ENO1/eno1::P<sub>ENO1</sub>-yGFP3* | This study |
| Δ/Δ ume6 | EGFP | *ade2::hisG/ade2::hisG ura3::imm434/ura3::imm434 ume6Δ::URA3/ume6Δ::NAT1 ENO1/eno1::P<sub>ENO1</sub>-yGFP3* | This study |
| Δ/+ ume6 (URA3) | EGFP | *ade2::hisG/ade2::hisG ura3::imm434/ura3::imm434 ume6Δ::URA3/UME6, ENO1/eno1::P<sub>ENO1</sub>-yGFP3* | This study |
| Δ/+ ume6 (NAT1) | SC5314 | *ume6Δ::NAT1/UME6* | This study |
| *C. neoformans* |  |  |  |
| H99 |  | Serotype A, clinical isolate | （62） |
| *A. fumigatus* |  |  |  |
| NBRC33022 |  | Reference strain | NBRC |
| *S. cerevisiae* |  |  |  |
| X2180-1B |  | Laboratory strain | (63) |
| Σ1278b |  | Filamentous laboratory strain | (64) |
| *S. pombe* |  |  |  |
| 972h<sup>–</sup> |  | Reference strain | （65） |

To estimate the number of viable fungal cells used for inoculation, the optical density at 600 nm (OD600) and colony-forming units (CFU) of each fungal culture were determined in multiple independent experiments, and calibration curves relating OD600 to CFU were generated. These calibration curves were used to estimate the CFU administered to frogs based on the OD600 of 24- or 48-h cultures used in each infection experiment.

*Aspergillus fumigatus* was cultured on potato sucrose agar (PSA) at 25°C for 96 h. Conidia were harvested using a cell scraper with PBS containing 0.2%Tween 20. The resulting suspension was passed through a cell strainer to remove large hyphal fragments, and the number of conidia was determined using a Bürker-Türk hemocytometer. The conidial suspension was centrifuged at 16,200 × *g* for 10 min, and the pellet was resuspended in PBS containing 0.2%Tween 20 and adjusted to the desired concentration.

Heat-killed fungal cells were prepared by centrifuging fungal cultures or conidial suspensions at 16,200 × g for 10 min. After removal of the supernatant, the fungal pellets were resuspended in PBS and autoclaved at 121°C for 20 min.

### Antifungal agents

The doses of antifungal agents were determined based on an average frog body weight of 2.5 g and were adjusted to approximate the body weight-based doses used clinically in pediatric patients. Amphotericin B was dissolved in dimethyl sulfoxide (DMSO) and diluted with PBS to a final concentration of 100 μg/mL. Micafungin sodium (Sawai Pharmaceutical Co., Ltd., Japan) was dissolved in 0.9%NaCl to a final concentration of 300 μg/mL. Antifungal solutions were stored at −30°C and thawed before use.

### Intraperitoneal inoculation

Fungal suspensions were drawn into 1-mL tuberculin syringes (Terumo, Tokyo, Japan) fitted with 27-gauge, 3/4-inch needles. Frogs were anesthetized by cooling on ice, and 50 μL of each fungal suspension was injected intraperitoneally through the abdominal wall above the left hind limb. Five frogs were used per group in each experiment. Following inoculation, frogs were maintained at 32°C in the dark in tanks containing 500 mL of water. Frogs recovered from cold anesthesia immediately after inoculation. PBS-injected frogs were included as negative controls in all experiments to control for potential effects of cold anesthesia and the injection procedure.

### Antifungal treatment

For immediate-treatment experiments, frogs were intraperitoneally inoculated with *C. albicans* ATCC 10231 as described above and immediately administered 50 μL of amphotericin B (5 μg/frog; approximately 2 mg/kg) or micafungin (15 μg/frog; approximately 6 mg/kg) intraperitoneally. The frogs were subsequently maintained at 32°C in water containing chloramphenicol (25 mg/L). Survival was recorded at 0, 12, 24, 36, 48, 72, and 96 h after treatment.

For delayed-treatment experiments, frogs were intraperitoneally inoculated with *C. albicans* ATCC 10231 and maintained at 32°C. At 6 h after inoculation, the frogs were again anesthetized by cooling on ice and administered 50 μL of amphotericin B (5 μg/frog; approximately 2 mg/kg) or micafungin (15 μg/frog; approximately 6 mg/kg) intraperitoneally. The frogs were subsequently maintained at 32°C in water containing chloramphenicol (25 mg/L), and survival was recorded at 12-h intervals after treatment.

For the corresponding vehicle controls, frogs were administered 50 μL of 0.2%DMSO in PBS for the amphotericin B experiments or 50 μL of 0.9%NaCl for the micafungin experiments. Vehicle-treated frogs were maintained under the same conditions as the antifungal-treated groups.

### Determination of LD50

For each experiment, three concentrations of fungal suspensions were prepared by serial dilution in PBS. Five frogs were intraperitoneally inoculated with each fungal dose, and survival was recorded at 72 h after inoculation. Each experiment was performed independently at least three times. Survival data were plotted against the administered fungal dose, and dose-response curves were fitted by logistic regression using R statistical software. The 50% lethal dose (LD50) and its standard error were estimated from the fitted logistic regression model.

### Determination of fungal burdens in blood and organs

Frogs were intraperitoneally inoculated with *C. albicans* ATCC 10231 and maintained at 32°C. At the indicated time points after inoculation, frogs were removed and anesthetized by immersion in 2 g/L Ethyl m-Aminobenzoate Methanesulfonate (MS-222; Nacalai Tesque, Kyoto, Japan). Blood was collected by severing the right hind limb at its base. The liver, kidneys, and brain were then excised, washed with PBS, and weighed. Each organ was homogenized in PBS using a Polytron homogenizer. Blood samples and organ homogenates were serially diluted 10-fold in 0.9%NaCl to generate seven-step dilution series. Five microliters of each dilution was spotted onto Sabouraud chloramphenicol agar plates and incubated at 30°C for 24 h. Colonies were counted as previously described (50). Fungal burdens were calculated based on colony counts and the corresponding dilution factors and expressed as CFU/mL of blood or CFU/g of tissue.

### RNA extraction from frog kidneys

Frogs were intraperitoneally inoculated with *C. albicans* ATCC 10231, *S. cerevisiae* X2180-1B, or PBS and maintained at 32°C. At 24 h after inoculation, frogs were anesthetized by immersion in 2 g/L MS-222. The kidneys were excised, immediately transferred to 2-mL tubes, and snap-frozen in liquid nitrogen.

Total RNA was extracted as described previously (51), with minor modifications. Briefly, frozen kidney tissues were homogenized in 600 μL of TRIzol reagent (Life Technologies Japan, Tokyo, Japan) using a Polytron homogenizer. The homogenates were centrifuged at 12,000 × *g* for 10 min at 4°C, and the supernatants were transferred to 1.5-mL tubes and incubated at room temperature for 5 min. Chloroform (120 μL) was added, and the samples were vortexed, incubated at room temperature for 3 min, and centrifuged at 12,000 × *g* for 15 min at 4°C. The aqueous phase (300 μL) was transferred to a new tube and mixed with 150 μL of a salt solution containing 0.8 M sodium citrate and 1.2 M NaCl. After addition of 150 μL of 2-propanol (Nacalai Tesque), the samples were vortexed, incubated at room temperature for 10 min, and centrifuged at 12,000 × *g* for 10 min at 4°C.

The supernatant was removed, and the RNA pellet was washed with 1 mL of 75%ethanol. After centrifugation at 7,500 × *g* for 5 min at 4°C, the ethanol was removed, and the pellet was air-dried for 10 min and dissolved in 30–50 μL of RNase-free water. Three independent experiments were performed, and RNA samples were obtained from a total of nine frogs per experimental group.

### RT-qPCR

For each sample, 2 μg of total RNA was adjusted to a volume of 16 μL with RNase-free water and treated with 2 μL of 10× RQ1 DNase reaction buffer and 2 μL of RQ1 RNase-Free DNase (Promega, Madison, WI, USA) at 37°C for 45 min. The reaction was terminated by adding 2 μL of RQ1 DNase Stop Solution, followed by incubation at 65°C for 10 min. cDNA was synthesized using the PrimeScript RT Reagent Kit (Perfect Real Time; Takara Bio, Shiga, Japan) according to the manufacturer’s instructions. Serial dilutions of the resulting cDNA were prepared, and quantitative PCR was performed using KOD SYBR qPCR Mix (Toyobo, Osaka, Japan), the appropriate primer pairs (Table 2), and a QuantStudio 3 Real-Time PCR System (Thermo Fisher Scientific, Waltham, MA, USA). Primers for *actb*, *tnf*, and *il1b* were described in previous studies using *X. laevis* (52–54). β-Actin was used as the internal reference. Relative transcript levels were calculated from standard curves generated for each primer pair and normalized to β-actin.

**Table 2.**
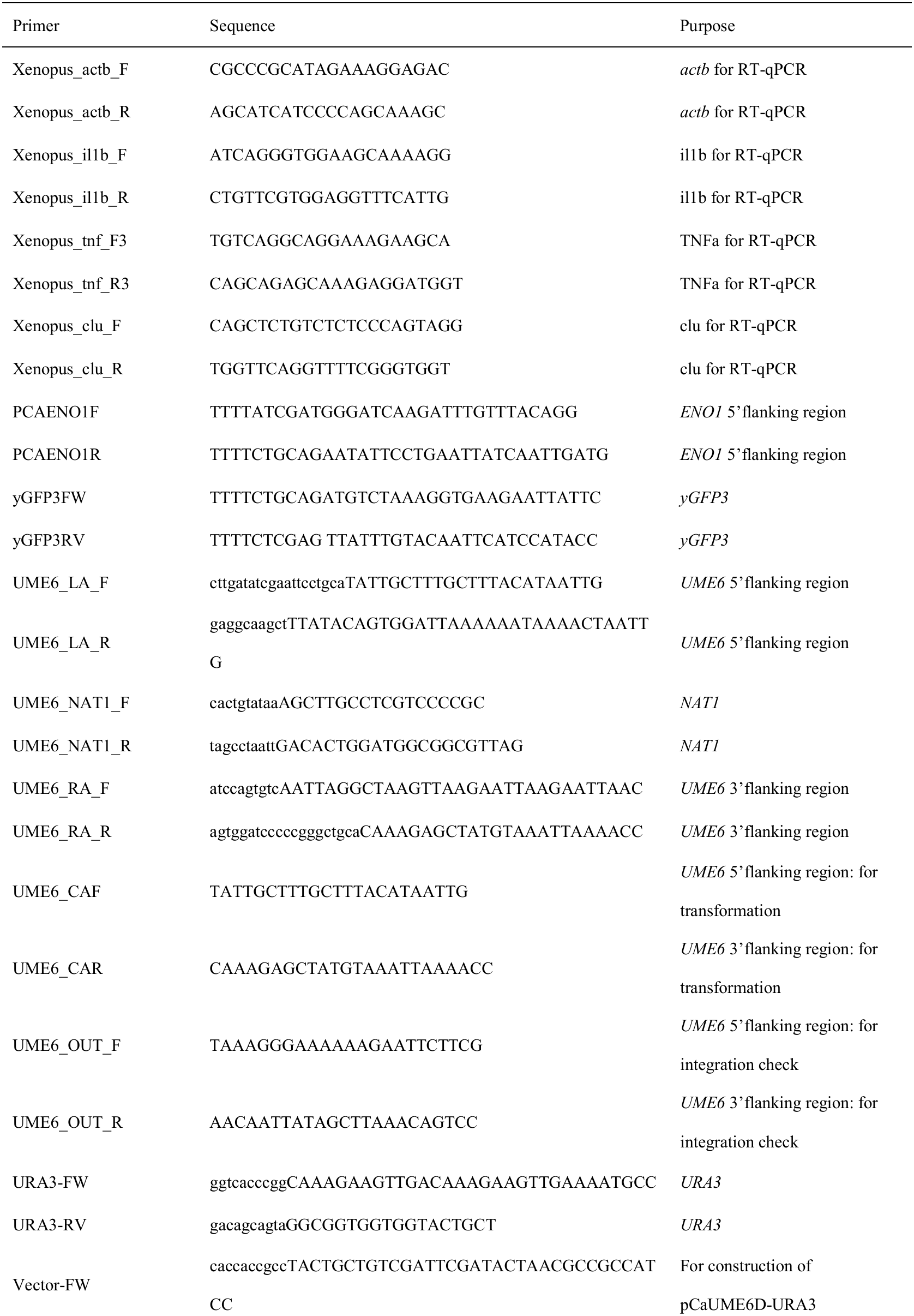
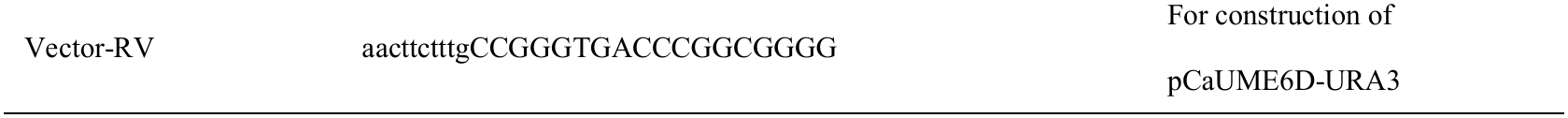
Oligonucleotide primers used in this study.

### Gross examination of the abdomen and kidneys

Frogs were intraperitoneally inoculated with *C. albicans* ATCC 10231, *S. cerevisiae* X2180-1B, or PBS and maintained at 32°C. At 24 h after inoculation, frogs were anesthetized by immersion in 2 g/L MS-222 and photographed. The abdominal cavity was then opened, and the internal organs were photographed. The kidneys were examined in situ using a Leica MZ10 F stereomicroscope (Leica Microsystems).

### Histological analysis

Frogs were intraperitoneally inoculated with *C. albicans* ATCC 10231, *C. albicans* SC5314, *S. cerevisiae* X2180-1B, or PBS and maintained at 32°C. At 24 h after inoculation, frogs were anesthetized by immersion in 2 g/L MS-222. The kidneys were excised and fixed in 10%neutral buffered formalin (NBF) for approximately 2 days. After fixation, the tissues were washed with Milli-Q water and dehydrated sequentially in 70%, 80%, 90%, 95%, and 100%ethanol for 30 min each, followed by an additional 30-min incubation in 100%ethanol. The tissues were cleared three times in xylene for 30 min each and infiltrated three times with paraffin at 60°C for 1 h each. The kidneys were then embedded in paraffin. Paraffin-embedded tissues were sectioned at a thickness of 5 μm using a Leica SM2000 R microtome (Leica Microsystems). Sections were mounted on glass slides and dried at 37°C for approximately 18 h.

For hematoxylin and eosin (H&E) staining, sections were deparaffinized twice in xylene for 10 min each and rehydrated by sequential incubation twice in 100%ethanol for 5 min each and once each in 95% ethanol, 70%ethanol, and water for 5 min. Sections were stained with hematoxylin solution (FUJIFILM Wako) for 10 min, washed with water, and incubated in water for 10 min. Sections were then stained with eosin Y solution (FUJIFILM Wako) for 5 min and washed with water. The sections were dehydrated in 70%ethanol for 5 min, followed by two 5-min incubations each in 95% and 100%ethanol. After two 5-min incubations in xylene, the sections were mounted with Entellan New (Merck, Darmstadt, Germany).

For periodic acid-Schiff (PAS) staining, sections were deparaffinized and rehydrated as described above. PAS staining was performed using a PAS staining kit (Muto Pure Chemicals, Tokyo, Japan) according to the manufacturer’s instructions. Briefly, sections were treated with periodic acid for 10 min and washed twice with water for 5 min. Schiff reagent was applied, and the sections were incubated at 37°C for 30 min. The sections were then treated twice with sulfurous acid solution for 5 min and washed with water. Sections were counterstained with Mayer’s hematoxylin for 15 min, washed with water, and incubated in a 60°C water bath for 10 min to develop the color. The sections were subsequently dehydrated, cleared, and mounted as described for H&E staining. H&E- and PAS-stained sections were examined using a ZEISS Axiovert 5 microscope (Carl Zeiss Microscopy) equipped with 10× and 63× objective lenses.

### Plasmid construction

The plasmid pCAIEGFP3 was constructed as follows. A fragment containing the *ENO1* promoter region (nucleotides −528 to −6) was amplified by PCR using primers PCAENO1F and PCAENO1R and cloned into the *Cla*I/*Pst*I sites of pCRW3 (55). The yGFP3 sequence was amplified from pYGFP3 (56) using primers yGFP3FW and yGFP3RV and cloned into the *Pst*I/*Xho*I sites of the resulting plasmid to generate pCAIEGFP3.

The plasmid pCaUME6D-NAT was constructed by assembling three DNA fragments into *Pst*I-digested pBluescript SK II(+): the 5′-flanking region of *UME6* (nucleotides −400 to −1), a 1,212-bp fragment containing the *NAT1* gene together with the *MET3* promoter from pNAT (57), and the 3′-flanking region of *UME6* (nucleotides +2554 to +2954). The fragments were assembled using the NEBuilder DNA Assembly Kit (New England Biolabs).

To construct pCaUME6D-URA3, the *NAT1*-containing fragment of pCaUME6D-NAT was replaced with a fragment containing the *URA3* coding region together with approximately 1-kb upstream and downstream flanking regions using the NEBuilder DNA Assembly Kit. The *URA3*-containing fragment and the vector backbone were amplified using primer pairs URA3-FW/URA3-RV and Vector-FW/Vector-RV, respectively.

Unless otherwise indicated, DNA fragments were amplified from *C. albicans* SC5314 genomic DNA, *S. cerevisiae* W303 genomic DNA, or the indicated plasmids, as appropriate. Primer sequences are listed in Table 2.

### Construction of *C. albicans* gene deletion mutants

Strain EGFP was generated by integrating AccI-linearized pCAIEGFP3 into the ENO1 locus of strain CAI8 using the lithium acetate transformation method, as described previously (58, 59). Subsequent gene disruption and complementation strains derived from EGFP were generated by electroporation.

Electrocompetent *C. albicans* cells were prepared using a protocol modified from that provided by NEPA GENE Co., Ltd. Cells were grown overnight in YPD medium (1% yeast extract, 2% Bacto Peptone, and 2% glucose) at 30°C. An aliquot of the overnight culture was transferred to fresh YPD medium and incubated at 30°C with shaking at 180 rpm until the cell density reached approximately 2 × 10⁷ cells/mL (OD600 ≈ 1.0). Cells were harvested by centrifugation at 8,000 × g for 3 min at room temperature and resuspended in 10 mL of HEPES-supplemented YPD medium. After the addition of 1 mL of TE buffer and 1 mL of 1 M lithium acetate, the suspension was incubated at 30°C with shaking at 100 rpm for 30 min. Dithiothreitol (DTT; 250 μL of a 1 M solution) was then added, and the suspension was incubated for an additional 10 min under the same conditions. The cells were washed repeatedly with ice-cold 10% glycerol until the electrical resistance of the suspension was within the appropriate range and were finally resuspended in approximately 800 μL of ice-cold 10% glycerol. The resulting electrocompetent cell suspension contained approximately 7 × 10⁸ cells/mL. Aliquots of 100 μL were stored at −80°C until use.

For electroporation, frozen electrocompetent cells were rapidly thawed, and 40 μL of the cell suspension was mixed with 40 μL of 1 M sorbitol containing 2–3 μg of DNA. After incubation on ice for 5 min, the mixture was transferred to a prechilled electroporation cuvette with a 1-mm gap and incubated on ice for an additional 5 min. Electroporation was performed using an N.Porator electroporator (NEPA GENE Co., Ltd.) at 1.5 kV. The sample impedance immediately before electroporation was typically 2–4 kΩ. Immediately after pulsing, 1 mL of prewarmed 1 M sorbitol was added for URA3 selection, whereas 1.5 mL of prewarmed YPD medium was added for nourseothricin or hygromycin selection. Cells were allowed to recover at 30°C with shaking at 250 rpm for 3 h for URA3 selection, 14 h for nourseothricin selection, or 6 h for hygromycin selection. The cells were then plated on the appropriate selective medium and incubated at 37°C for 1–2 days.

To generate the heterozygous UME6 deletion strain Δ/+ ume6 (URA3), one UME6 allele in strain EGFP was replaced with URA3 using a disruption fragment amplified from pCaUME6D-URA3 with primers UME6_CAF and UME6_CAR. To generate the homozygous UME6 deletion strain Δ/Δ ume6, the remaining UME6 allele in Δ/+ ume6 (URA3) was replaced with NAT1 using a disruption fragment amplified from pCaUME6D-NAT with the same primer pair.

PCR amplification of the DNA fragments used to construct the UME6 deletion was performed using KAPA HiFi DNA polymerase (NIPPON Genetics). Correct integration of the transforming DNA at the respective target loci was confirmed by PCR for all strains (Table 2).

### Statistical analysis

Statistical analyses were performed using GraphPad Prism (GraphPad Software, Boston, MA, USA) or R for Windows, as indicated below. Kaplan-Meier survival curves from the antifungal treatment experiments were compared using the log-rank (Mantel-Cox) test. For experiments evaluating fungal virulence genes, Kaplan-Meier survival curves were compared using the log-rank test with Holm-Šídák correction for multiple comparisons.

Dose-response curves were generated by logistic regression in R for Windows by modeling survival as a function of the fungal dose administered to each frog. The 50% lethal dose (LD50) and its standard error were estimated using the glm function in R. Differences between two dose-response curves were evaluated by likelihood ratio tests using the anova function. The R code used for these analyses is available in a GitHub repository and has been archived in Zenodo (doi:10.5281/zenodo.14978657).

Differences in fungal burdens in organs were analyzed using Dunnett’s multiple-comparisons test in GraphPad Prism. Differences in gene expression determined by RT-qPCR were analyzed using unpaired *t* tests with Welch’s correction. A *P* value of <0.05 was considered statistically significant.

### Ethics approval

This study was conducted in strict accordance with the Fundamental Guidelines for Proper Conduct of Animal Experiments and Related Activities in Academic Research Institutions under the jurisdiction of the Ministry of Education, Culture, Sports, Science and Technology of Japan (2006). All procedures involving frogs were conducted in accordance with the Regulations for Animal Care and Use of Okayama University and were approved by the Animal Use Committee of Okayama University (approval no. OKU-2025603).

## Acknowledgements

This study was supported by the Japan Society for the Promotion of Science (JSPS) Grants-in-Aid for Scientific Research (Grants 26K23680, 26K02250, 24K01760, 24K21872). This work was also supported by the Institute for Fermentation, Osaka (IFO) (G-2026-2-010) and the Association for Research on Lactic Acid Bacteria. We thank Dr. Taiga Miyazaki for providing pNAT, the Biological Resource Center, National Institute of Technology and Evaluation (NBRC) for providing the microbial strains used in this study.

